# A survey of multi-targeting and off-targeting sgRNAs across five genome-wide CRISPR-Cas9 knockout sgRNA libraries with GuideRefine

**DOI:** 10.64898/2026.08.20.746015

**Authors:** Stefanus Bernard, Michael D. Rainey, Corrado Santocanale, Colm J. Ryan

## Abstract

Pooled genome-wide CRISPR-Cas9 knockout (CRISPR-KO) screening is a powerful approach for discovering new biology and identifying genetic vulnerabilities in cancers. This approach uses the Cas9 nuclease in combination with sgRNA libraries, typically consisting of 4-8 sgRNAs to induce mutations in each target gene. A critical assumption is that the effect of each sgRNA is solely due to Cas9 editing of the target gene. However, libraries can contain sgRNAs that direct Cas9 to multiple locations, thus potentially introducing bias into gene hit lists and leading to flawed biological hypotheses.

Here we have developed GuideRefine, a pipeline to detect multi-targeting and off-targeting sgRNAs. GuideRefine outputs a virtual refined sub-library containing only on-target sgRNAs. Using GuideRefine with T2T-CHM13 as the reference genome, we surveyed the Brunello, TKOv3, Yusa, Avana, and Jacquere libraries, finding that ∼7.5% to ∼16% of sgRNAs are potentially problematic. We confirmed that multi-targeting sgRNAs disproportionately impair cell fitness and that sgRNAs aligning to more than one location with a single mismatch can also reduce fitness, although to a lesser extent.

After flagging problematic sgRNAs and creating virtual “on-target only” sub-libraries, ∼10% to ∼16% of genes lose critical representation (< 3 sgRNAs per gene). Intriguingly, a set of 467 genes, characterised by short CDS length and lower PAM site density, have fewer than three sgRNAs in all sub-libraries, suggesting they cannot be well-targeted using current libraries.

We anticipate that GuideRefine, together with caution in assessing the effects of problematic sgRNAs, will help prioritise biologically relevant hits.

## INTRODUCTION

Pooled genome-wide CRISPR-Cas9 knockout (CRISPR-KO) screens are an experimental approach that systematically disrupt thousands of genes across a cell population to identify their associated phenotypes (Wang et al. 2014). This technology has been widely used to reveal novel genetic vulnerabilities in cancer cell lines and to identify genes whose loss confers resistance or sensitivity to specific drugs (Behan et al. 2019; Pacini et al. 2024). The approach relies on the Cas9 nuclease cleaving the target DNA at genomic sites specified by single guide RNAs (sgRNAs) delivered as pooled libraries to induce loss-of-function mutations in target genes (Jinek et al. 2012; Wang et al. 2014). Each sgRNA contains a spacer sequence, typically 20 nucleotides sequence, that directs Cas9 to a complementary genomic target located next to a Protospacer Adjacent Motif (PAM) site, upon which Cas9 induces double-strand breaks that are mostly repaired by the non-homologous end joining pathway, leading to loss-of-function mutations in most situations (Jinek et al. 2012; Cong et al. 2013). To ensure robust gene-level statistical inference, and to account for variability in sgRNA targeting efficiency, a typical sgRNA library contains 4 to 8 sgRNAs per protein-coding gene (Otten and Sun 2020). CRISPR-KO screen analysis then aggregates the effects of all sgRNAs targeting a given gene to generate a ranked list of genes impacting cell fitness or sensitivity to treatments (Zhao et al. 2022). A critical assumption of this approach is that each sgRNA can be connected to a single gene.

Several CRISPR-KO sgRNA libraries have been developed with a varying total number of sgRNAs and target genes. Widely used CRISPR-KO libraries contain 70,000 – 90,000 sgRNAs targeting approximately 19,000 protein-coding genes, with an average of 4-5 sgRNAs per gene (Doench et al. 2016; Tzelepis et al. 2016; Hart et al. 2017; Meyers et al. 2017; Drepanos et al. 2026). These libraries are typically designed using computational algorithms to optimise sgRNA on-target activity while minimising possible off-target cleavage (Jia et al. 2025). For instance, the Brunello library applied the Rule Set 2, an on-target activity scoring algorithm that ranks sgRNAs by predicted Cas9 cutting activity based on spacer nucleotide composition, alongside the Cutting Frequency Determination (CFD) score, an off-target scoring metric weighted by nucleotide position and mismatch type, to minimise off-target effects (Doench et al. 2016). Similarly, Toronto Knockout version 3 (TKOv3) library selects high-performing sgRNAs based on SeqScore, a score that predicts Cas9 cleavage activity based on the identity of nucleotides at each position of the sgRNA spacer sequence, derived from sgRNA performance in CRISPR-KO screens (Hart et al., 2017).

A known limitation in CRISPR-KO screens is the short length of sgRNA spacer sequences that help guide Cas9, 20 nucleotides for most libraries and 19 nucleotides sequence for the Yusa (Project Score) library. This means that identical or highly similar sequences can occur elsewhere in the genome. Thus, libraries can contain sgRNAs that direct Cas9 to multiple genomic locations due to perfect alignments (multi-target sgRNAs) (Aguirre et al. 2016; Munoz et al. 2016; Fortin et al. 2019), or closely aligned sequences due to the single or double nucleotide mismatch tolerance between the Cas9-sgRNA complex and DNA sequence during target binding (Fu et al. 2013). The presence of these problematic sgRNAs can bias the gene hit lists from CRISPR-KO screen analyses, and potentially lead to erroneous formulation of biological hypotheses (Fortin et al. 2019). To mitigate this potential confounding factor, library quality control can be performed by mapping all sgRNA sequences to the reference genome, followed by computational removal of suspected problematic sgRNAs from the library. However, this remains an underutilised practice, with only a limited number of studies implementing such *in-silico* filtering of problematic sgRNAs (Tycko et al. 2019; Kim and Hart 2021; Rainey et al. 2025). We have undertaken this approach in our own chemogenetic pooled CRISPR-KO screens, aimed at identifying genes modulating sensitivity to CDC7 inhibitors. We used the Cellecta sgRNA library, containing approximately 150,000 sgRNAs targeting 19,000 protein-coding genes with 8 sgRNAs per gene. As one of the earliest commercial sgRNA libraries, it may contain problematic sgRNAs, and we developed a ‘library cleaning’ tool to refine this library prior to downstream analysis (Rainey et al. 2025). While this approach is an essential quality control step, in practice, researchers must define their own filtering parameters and construct their own pipeline. Currently, there is no dedicated pipeline available that can refine an sgRNA library by identifying problematic sgRNAs.

*In-silico* removal of problematic sgRNAs inevitably decreases the representation of sgRNAs targeting specific genes (Doench et al. 2016) and consequently, some genes may lose sgRNA coverage as they fall below the minimum coverage threshold for reliable statistical gene level inference (typically of three or more sgRNAs per gene) (Kim and Hart 2021; Zhao et al. 2022). An sgRNA library designed with near-perfect specificity could theoretically reduce the need for *in-silico* removal of problematic sgRNAs, but it is not clear to what extent current libraries reflect this ideal-case. An up-to-date systematic survey of multi-targeting and off-targeting sgRNAs across existing libraries may serve as a valuable reference to identify limitations with existing libraries and to improve future library design.

In this study, we address both gaps by further developing our ‘library cleaning’ tool into GuideRefine, an R-based computational pipeline for *in-silico* detection and removal of multi-targeting and off-targeting sgRNAs from CRISPR-KO libraries. Currently available tools such as crisprVerse provide a strategy to annotate and predict off-target activity for all sgRNAs in existing libraries, but do not computationally remove problematic sgRNAs from the library (Hoberecht et al. 2022). Our pipeline addresses this by applying a computational detection and removal of problematic sgRNAs based on a predefined criterion of problematic sgRNAs. GuideRefine outputs several files including ready-to-use refined *in-silico* library for downstream analysis of CRISPR-KO screens, a list of problematic sgRNAs, an HTML summary report after library refinement, and an excel file containing the sgRNA representation for each gene at each refinement step.

By applying GuideRefine, we present an updated survey of multi-targeting and off-targeting sgRNAs across five major CRISPR-KO libraries, including the Brunello (Sanson et al. 2018), TKOv3 (Hart et al. 2017), Yusa (Project Score) (Tzelepis et al. 2016), Avana (Broad Dependency Map) (Meyers et al. 2017), and Jacquere libraries (Drepanos et al. 2026). We provide the first systematic cross-library comparison of problematic sgRNAs using consistent filtering criteria. This survey was also performed due to the release of T2T-CHM13 reference genome, that currently represents the complete assembly of the human genome (Nurk et al. 2022), thus enabling the identification of problematic sgRNAs that were potentially missed in the previous GRCh38 genome build. For each library, GuideRefine produces an *in-silico* refined version that only contains the subset of sgRNAs predicted to have a single on-target effect. We provide the resulting *in-silico* refined libraries as a resource for broader research community who use any of these five libraries in their CRISPR-KO screens, enabling immediate use in downstream analysis without requiring further quality control steps.

## MATERIALS AND METHODS

### Brunello library and sgRNA log-fold change data

Brunello sgRNA library sequences were obtained from the supplementary data of Sanson et al (2018). The Brunello library is composed of 77,441 sgRNAs targeting 19,115 genes, with an average of four sgRNAs/gene. This library also contains 1,000 non-targeting control sgRNAs. The sgRNA read count data was retrieved from the same study, in which a CRISPR-KO screen was performed in A375 cells using the Brunello library with modified trans-activating CRISPR RNA (tracrRNA).

The sgRNA level normalisation and log-fold change calculation was performed similarly as described by Fortin et al (2019). First, we filtered out all sgRNAs with fewer than 30 reads in the plasmid DNA (pDNA) samples. Read counts for each replicate were then log-transformed, and log-fold changes were computed relative to the pDNA. Log-fold change values were subsequently normalised by centering around the median of non-targeting control sgRNAs. Log-fold change values across replicates were scaled by the absolute mean log-fold change of sgRNAs targeting 217 constitutive core essential genes. The log-fold change values were then averaged across replicates to obtain a representative sgRNA-level activity score for each sgRNA.

### TKOv3 library and sgRNA log-fold change data

The Human Toronto Knockout version 3 (TKOv3) sgRNA library sequence file was retrieved from AddGene (https://www.addgene.org/pooled-library/moffat-crispr-knockout-tkov3/). This library consists of 71,090 sgRNAs targeting 18,056 genes. TKOv3 library also contains 142 control sgRNAs targeting EGFP, LacZ, and luciferase. The sgRNA read count data was obtained from Colic et al (2019), in which a CRISPR-KO screen was performed in RPE1 cells using the TKOv3 library (Colic et al. 2019). The read count data used in this study are publicly available at https://figshare.com/articles/dataset/Readcounts/8799215?file=16170896 (matrix-reads-by-gRNA-RPE1-drugZ.txt). We restricted our analysis to read counts representing cell proliferation only (RPE1_T0, RPE1_T3A_CTRL, RPE1_T3B_CTRL). The read count filtering, log-fold change computation, scaling of constitutive core essential genes, and data normalisation were performed following the same approach applied to the Brunello library.

### Yusa (Project Score) library and sgRNA log-fold change data

Yusa (Project Score) sgRNA library sequences were retrieved from AddGene (https://www.addgene.org/pooled-library/yusa-crispr-knockout-human-v1/). This library comprises 90,709 sgRNAs targeting 18,009 genes, with an average of five sgRNAs/gene, where each sgRNA consists of 19-nucleotide sequence rather than the canonical 20 nucleotides. sgRNA-level raw log-fold change data was obtained from DepMap release 25Q2 (KYLogfoldChange.csv) that represents sgRNA activity across 316 cell lines screened by using Yusa library. Unlike Brunello and TKOv3 libraries, the Yusa data retrieved from DepMap are already provided as log-fold changes and therefore do not require read count filtering or log-transformation prior to analysis. The sgRNA log-fold change data values were subsequently normalised using a similar approach that described by Fortin et al (2019) for the Avana library. Briefly, sgRNA log-fold change values across all replicates were centered using their median absolute deviation. The log-fold change values for sgRNAs targeting constitutive core essential genes (217 genes), as defined by Hart et al (2014), were further scaled by the absolute mean log-fold change across all cell lines (Hart et al. 2014). Finally, the log-fold change values were averaged across all cell lines and replicates to obtain a representative log-fold change value for each of the sgRNA in Yusa (Project Score) library. Unlike Fortin et al (2019), the copy number correction was not applied in this study as our analysis focuses on sgRNA-level activity aggregated across all cell lines without investigating a specific cell line.

### Avana library and sgRNA log-fold change data

The Avana sgRNA library sequences were obtained from the supplementary data of Meyers et al. (2017) and comprises of 71,826 sgRNAs targeting 17,487 protein-coding genes, with an average of four sgRNAs/gene. This library also contains 995 non-targeting control sgRNAs. The sgRNA log-fold change data for Avana library were obtained from The Cancer Dependency Map (DepMap) project release 25Q2 (AvanaLogfoldChange.csv). This DepMap release 25Q2 has 1,116 unique cell lines across many cancer subtypes. Log-fold change data normalisation and scaling of constitutive core essential genes were performed following the same approach applied to the Yusa (Project Score) library.

### Jacquere library and sgRNA log-fold change data

The Jacquere sgRNA library sequences and corresponding sgRNA log-fold change data were obtained from Drepanos et al. (2026), in which a CRISPR-KO screen was performed in A375 cells using the Jacquere library. We noted that Drepanos et al. analysed a subset of the Jacquere library that contains an average of three sgRNAs per gene (Drepanos et al. 2026). However, we chose to analyse the full Jacquere library with four sgRNAs per gene, as provided in the Drepanos et al. supplementary file to maintain consistency with the majority of libraries in our survey.

We obtained the Jacquere sgRNA library sequences file from the GPP-Jacquere GitHub repository (Jacquere_PerGuideAnnotations_Quota4.csv, 80,124 sgRNAs). As the GuideRefine pipeline performs a gene symbol to exon annotation validation step, any sgRNAs with missing gene symbol have no annotation available for validation and were therefore excluded prior to downstream analysis. This resulted in a library of 78,784 sgRNAs targeting 19,735 genes, with an average of four sgRNAs per gene. This library also contains 1,000 intergenic controls sgRNA (ONE_SITE_INTERGENIC) and 1,000 non-targeting control sgRNAs (NO_SITE). sgRNA log-fold change data were acquired from the GPP-Jacquere GitHub repository (jacquereA375lfc.csv). As Drepanos et al. performed CRISPR-KO screen using three sgRNAs per gene rather than the full four, the log-fold change data were not available for all sgRNAs in the library. Therefore, we restricted our analysis to sgRNAs with available log-fold change data. sgRNA log-fold change was normalised by centering all log-fold change value using the median of all non-targeting control sgRNAs (NO_SITE). We then performed the scaling of sgRNA targeting constitutive core essential genes by using the absolute mean log-fold change of all sgRNAs and averaged log-fold change across replicates.

### Restricting library for eligible sgRNAs and genes

To ensure consistency in the gene universe across all five libraries, sgRNAs were filtered to retain only those targeting eligible protein-coding genes. Eligible protein-coding genes were defined by two criteria: (1) genes categorised as protein-coding by the HUGO Gene Nomenclature Committee (HGNC) (Seal et al. 2023), and (2) presence of a canonical mRNA transcript containing a CDS (Coding DNA Sequence) in the T2T-CHM13 annotation data. To obtain the gene sets representing the first criterion, the HGNC complete dataset (TSV format) was obtained from the HGNC download files portal (https://www.genenames.org/download/). Genes were retained if the locus_group field was annotated as “protein-coding gene” and the locus_type field as “gene with protein product”.

To retrieve the gene sets fulfilling the second criterion, the T2T-CHM13v2.0 GFF file and assembly report were obtained from the NCBI file transfer protocol server (https://ftp.ncbi.nlm.nih.gov/genomes/all/GCF/009/914/755/GCF_009914755.1_T2T-CHM13v2.0/). Canonical mRNA transcripts and CDS were extracted from the GFF file and converted into a CCDS (Consensus Coding Sequence)-like format containing exon coordinates for each gene. The assembly report was used to match Refseq accession IDs to their corresponding chromosomes. Data were further processed using *Tidyverse* package (Wickham et al. 2019), yielding a final table comprising of chromosome ID, gene symbol, strand, CDS start and end coordinates, and range of CDS. This annotation is formatted analogously to the CCDS standard (Pujar et al. 2018), and can also be used directly as GuideRefine input.

The intersection of gene symbols from the filtered HGNC data and the T2T-CHM13 CCDS-style annotation data yielded the final set of eligible protein-coding genes for subsequent analysis. Gene symbols for sgRNAs in all five libraries were standardised against current HGNC approved symbols using *HGNChelper*, an R package for identifying and updating outdated gene symbols (Oh et al. 2022). Libraries were then filtered to retain only sgRNAs targeting genes that fulfilled both criteria described above.

### Forge T2T-CHM13 BSgenome package in R

The Telomere-to-Telomere (T2T-CHM13) reference genome represents the first complete sequence of a human genome with a total of 3.055 billion base pairs (Nurk et al. 2022). Bioconductor provides the BSgenome assembly for T2T-CHM13 reference genome in GenBank format (accession ID: GCA_009914755.4). However, this format does not have the Gene Feature Format (GFF) annotation data that is required for GuideRefine input. To ensure compatibility and consistency in our analysis, we forged our own T2T-CHM13 reference genome in NCBI RefSeq assembly format (accession ID: GCF_009914755.1). We used *forgeBSgenomeDataPkgFromNCBI* function from the *BSgenomeForge* version 1.10.2 in R. This function retrieves the T2T-CHM13v2.0 reference genome fasta file in NCBI Refseq assembly format and forge it into a BSgenome-compatible data package, which enable the user to import it as a library in the R environment.

### sgRNA alignment classification

To facilitate the systematic survey of the problematic sgRNAs across all five libraries, we developed a custom R script to categorise each sgRNA based on our pre-defined criteria. The script takes as input the original library file, the sgRNA alignment file (intermediate output from GuideRefine), and a tabular file listing all computationally removed sgRNAs (final output from GuideRefine). This script outputs the flagged problematic sgRNAs and non-problematic sgRNAs in five libraries, provided as Supplementary Data, which can be integrated with the corresponding sgRNA log-fold change data. This combined dataset enabled the investigation of whether multi-targeting and single-mismatch sgRNAs across all five libraries disproportionately impair cell fitness.

### Comparison of problematic sgRNAs identified by using GRCh38 and T2T-CHM13

We compared problematic sgRNAs identified using GRCh38 and T2T-CHM13 by applying GuideRefine to all five libraries with both reference genome. The GRCh38 reference genome (Schneider et al. 2017), also known as hg38 in UCSC genome browser, was obtained from the UCSC genome browser portal (https://hgdownload.soe.ucsc.edu/goldenPath/hg38/bigZips/hg38.fa.gz). The T2T-CHM13v2.0 reference genome was obtained programmatically as described in the previous section (Forge T2T-CHM13 BSgenome package in R). All sgRNA sequences were aligned to each reference genome using *crisprBowtie* with a maximum tolerance of two mismatches (Langmead et al. 2009; Hoberecht et al. 2022). We then used the custom R script described in the previous section (sgRNA alignment classification) to flag all problematic and non-problematic sgRNAs. The proportion of problematic sgRNAs in all five libraries was then calculated and compared for both genome builds.

### Analysis of the problematic sgRNAs in Jacquere library identified by using GuideRefine and Drepanos et al. (2026)

To compare problematic sgRNAs in the Jacquere library identified by Drepanos et al. with those identified using GuideRefine, we obtained the Jacquere library per-guide annotation file (Jacquere_PerGuideAnnotations_Quota4.csv) from the GPP-Jacquere GitHub repository. Drepanos et al. flagged 2,024 sgRNAs in the Jacquere library with a CFD score of 1.0, or 100% CFD probability off-target sites (OTS) (Drepanos et al. 2026). To ensure consistency with the analysis implemented by Drepanos et al., the library refinement of the Jacquere library was performed using GuideRefine with the GRCh38 reference genome. The problematic sgRNAs identified by using the two approaches were visualised in the form of UpSet plot by using UpSetR version 1.4.0 (Conway et al. 2017).

As we are interested in analysing single mismatch sgRNAs exclusively identified by GuideRefine, we restricted our analysis where log-fold change data is available (see Jacquere library and sgRNA log-fold change data section). From a total of 6,807 single mismatch sgRNAs identified exclusively by GuideRefine, but not by Drepanos et al., only 5,204 single mismatch sgRNAs has an available log-fold change data. Off-targeting activity of sgRNAs in this set was scored using *crisprScore* version 1.14.0 implementing CFD scoring (Hoberecht et al. 2022). For sgRNAs with more than one predicted off-target alignment, we report the maximum CFD score across all off-target alignments per sgRNA, representing the alignment with the highest predicted off-target Cas9 cleavage activity.

### Retrieving total CDS length for each gene

We used the same T2T-CHM13v2.0 GFF file as described in the previous section (Restricting library for eligible sgRNAs and genes). A Txdb sqlite file was created for T2T-CHM13v2.0 GFF data by using *makeTxDbFromGFF* function in *GenomicFeatures* version 1.24.4 (Lawrence et al. 2013). CDS lengths were obtained for 467 genes with fewer than three sgRNAs and 18,507 other protein-coding genes by first subsetting the T2T-CHM13v2.0 genomic ranges data for each respective gene group. Within each group, the CDS regions were extracted using the *cdsBy* function. We filtered out overlapping CDS regions within each gene by using *reduce* function to eliminate redundancy. The length of non-redundant CDS per gene were then summed to yield the actual CDS length by base pairs for each gene. Both genes group were included in the analysis of CDS length and the number of possible protospacer sequences with an NGG PAM site in the CDS region.

### Generating spacer sequences in CDS region

All possible protospacer sites with an NGG PAM site within each CDS region were identified using *crisprDesign* version 1.12.0, a computational tool for spacer sequence design that is part of the larger *crisprVerse* ecosystem (Hoberecht et al. 2022). We used the *findSpacers* function to obtain a list of all possible spacer sequences targeting protospacers located in the target DNA sequence(s).

### sgRNAs aggregation and on-target activity scoring

Remaining sgRNAs targeting the 467 consistently underrepresented genes across four libraries (Brunello, TKOv3, Avana, and Jacquere) were aggregated into a composite of mini library. Yusa (Project Score) library was excluded from this strategy as its 19-nucleotide sgRNA sequences are not compatible with the 20-nucleotide sgRNAs of the other four libraries. Prior to on-target scoring, we filtered out sgRNAs targeting the same gene with identical sequence derived from multiple libraries to eliminate redundancy. sgRNA on-target activity was then scored by using *crisprScore* version 1.14.0 and implementing Rule Set 3, an on-target activity scoring framework that consider sgRNA sequence composition and small variations in the sequence of trans-activating CRISPR-RNA (tracRNA) (DeWeirdt et al. 2022). As the libraries included in this study were developed with different tracrRNA designs, Rule Set 3 scoring was conducted separately for each library group based on their original tracrRNA. Brunello, TKOv3, and Avana implemented the tracrRNA design from Hsu et al. (2013), while Jacquere was optimised using Chen et al. (2013) tracrRNA (Chen et al. 2013; Hsu et al. 2013). To construct a mini-composite library, the aggregated sgRNAs sequence were filtered by positive score (Rule Set 3 score > 0) to obtain high efficacy on-target sgRNAs and each gene should be covered by a minimum of three sgRNAs.

### Software and tools

Unless otherwise stated, analysis was performed in R version 4.5.2 (Ihaka and Gentleman 1996) using the *Tidyverse* version 2.0.0 (Wickham et al. 2019) and Bioconductor version 3.22 (Gentleman et al. 2004). sgRNA-level log-fold change normalisation was performed in Python version 3.12.7 using *Pandas* version 2.2.2 and *NumPy* version 1.26.4 (McKinney 2010; Harris et al. 2020).

### Statistical analysis

The non-parametric Wilcoxon rank-sum test was used to assess the statistical significance involving two group comparison and was performed using the *stats* package in R (Noether 1992). The Jonckheere-Terpstra trend test (Jonckheere 1954; Terpstra 1952), a non-parametric test to assess the monotonic trendline across ordered groups, was used to evaluate the association between the number of off-target alignment caused by multi-targeting and single-mismatch sgRNA to cell fitness, and was performed using *clinfun* package version 1.1.5 (Seshan and Whiting 2007).

### Large Language Models

Claude Sonnet version 4.6 (Anthropic), a Large Language Model (LLM), was used to assist in debugging and technical error correction in the GuideRefine pipeline, restructuring the GitHub repository for reader’s accessibility, and improving the grammatical clarity of this manuscript.

## RESULTS

### Development of the GuideRefine pipeline

To facilitate the computational detection and removal of multi-targeting sgRNAs and sgRNAs with high potential for off-targeting activity in CRISPR-KO screen analysis, we developed GuideRefine, an R-based computational pipeline for the systematic identification and filtering of problematic sgRNAs from CRISPR-KO libraries (**Figure 1a**). The pipeline was developed using R version 4.5.2 and Bioconductor version 3.22 (Ihaka and Gentleman 1996; Gentleman et al. 2004). This pipeline accepts the input of: (1) an sgRNA library in a tab-delimited text format, (2) reference genome annotation data, (3) a CDS annotation extracted from the reference genome, equivalent to the CCDS standard (Pujar et al. 2018), and (4) up-to-date HGNC annotation data (Seal et al. 2023). This pipeline is compatible with any reference genome available through Bioconductor *BSgenome* packages (Pagès 2017). The user can also provide their own reference genome by following the instructions described in *BSgenomeForge* (Pagès and Kakopo 2026). GuideRefine will align all sgRNA sequences to the input reference genome using *crisprBowtie* with a maximum of 2 mismatches allowed (Langmead et al. 2009; Hoberecht et al. 2022). The alignment process generates an intermediate file containing sgRNA alignments in csv format. This alignment file can be used to categorise each sgRNA and facilitates surveying problematic sgRNAs. Further detail is described in the sgRNA alignment classification section (see Methods).

**Figure 1.**
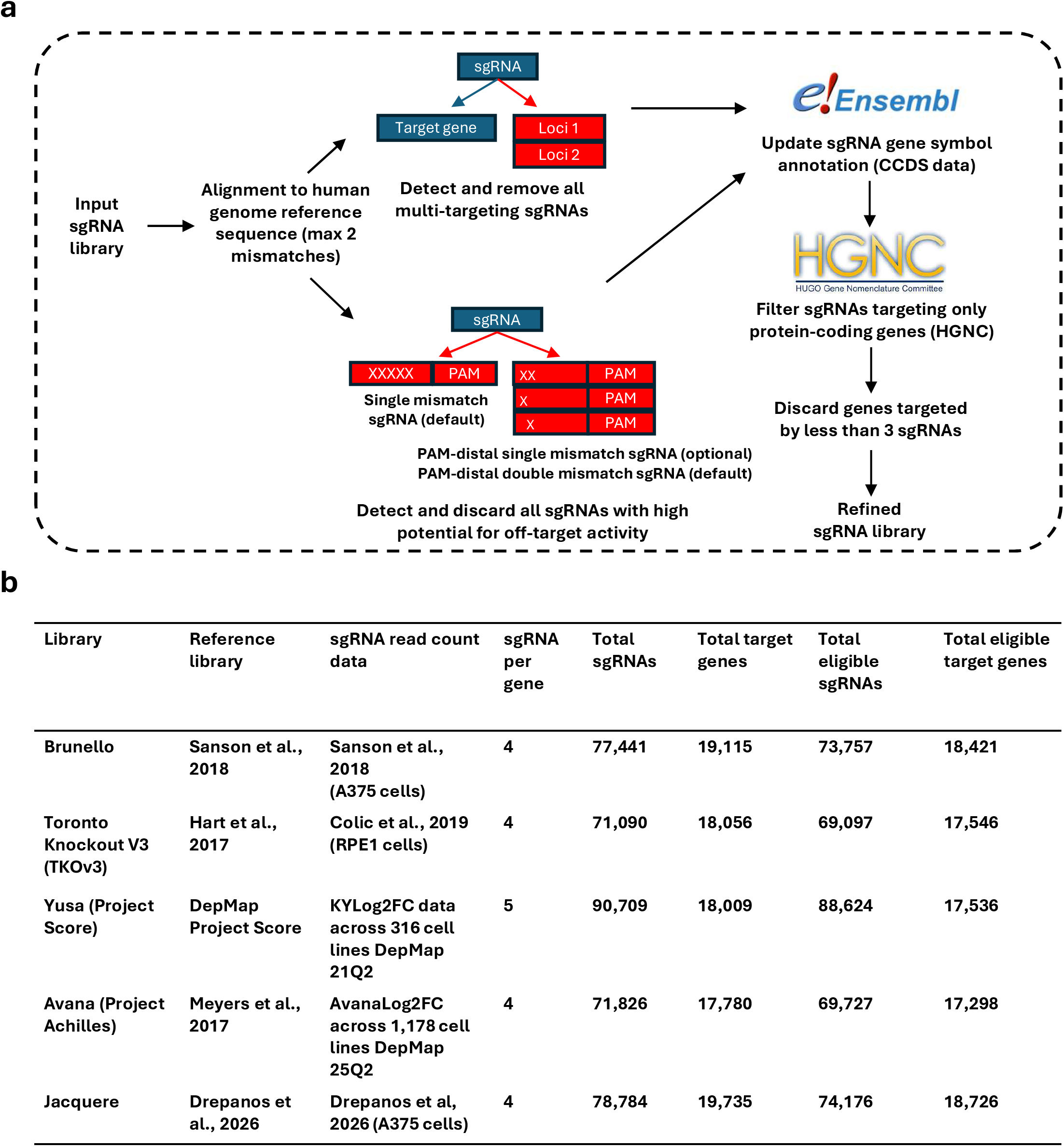
A survey of multi-targeting and off-targeting sgRNAs across CRISPR-KO libraries. **(a)** GuideRefine, an R-based pipeline for detecting and removing multi-targeting sgRNAs and sgRNAs with high potential for off-targeting activity in CRISPR-KO libraries. **(b)** Application of GuideRefine to survey multi-targeting sgRNAs and sgRNAs with high potential for off-targeting activity across five widely used CRISPR-KO libraries. The total eligible sgRNA and gene universe shown in panel **(b)** comprises of genes classified as protein-coding by HGNC and annotated with a canonical mRNA transcript present in the T2T-CHM13 GFF annotation data (see Methods).

The criteria for identifying problematic sgRNAs were adapted from DeKegel & Ryan (2019) and Fortin et al (2019) (De Kegel and Ryan 2019; Fortin et al. 2019). Multi-targeting sgRNAs are defined as sgRNAs that align perfectly to multiple protospacer location with PAM-sites in the genome. sgRNAs with high potential for off-targeting activity in this study are instead defined as: (1) all sgRNAs that align with a single mismatch in any location between the sgRNA spacer sequence and the target DNA, termed hereafter as single mismatch sgRNA; and (2) sgRNAs that align with single or double nucleotide mismatches at the two base positions located far from the PAM site, which were previously found to be associated with increased off-target activity (Fortin et al. 2019). By default, GuideRefine detects multi-targeting sgRNAs, single mismatch sgRNAs, and PAM-distal double mismatch sgRNAs. This stringency can be reduced by disabling the parameter to remove all single mismatch sgRNAs, which GuideRefine will then only filter out multi-targeting sgRNAs, PAM-distal single mismatch sgRNAs, and PAM-distal double mismatch sgRNAs as they are highly associated with increased off-target activity (Fortin et al. 2019).

GuideRefine will then use the input CDS data to annotate all sgRNAs based on their exon location within the target gene, verified against the genomic alignment coordinates. The negative control and intron control sgRNAs were excluded from this annotation step and were assessed only for multi-targeting and off-targeting activity. The assigning of sgRNAs to exons was performed using the *GenomicRanges* package in R (Lawrence et al. 2013). sgRNA that do not target an exon of any protein-coding gene as defined by HGNC were computationally removed from the library (Seal et al. 2023). The gene symbol corresponding to each sgRNA ID was updated based on the actual mapping between the sgRNA sequence and the target gene, to ensure the consistency with current HGNC approved symbols.

GuideRefine only retains all sgRNAs targeting protein-coding genes as defined by HGNC. To ensure sufficient representation of sgRNA number per gene, all genes covered by fewer than three sgRNAs (0-2 sgRNAs) after filtering were excluded from the sgRNA library. GuideRefine produces four final outputs: (1) a virtual sub-library containing only on-target sgRNAs with no predicted off-target activity; (2) an HTML report summarising the library cleaning process; (3) a tabular file detailing all computationally removed sgRNAs and the reasons for their elimination; and (4) an excel file that reports the sgRNA number of each gene upon each step-by-step filtering process.

### A survey of multi-target-/off-target sgRNAs across CRISPR-KO libraries

GuideRefine was used to survey multi-targeting and off-targeting sgRNAs across five major CRISPR-KO libraries: Brunello (Sanson et al. 2018), TKOv3 (Hart et al. 2017), Yusa (Project Score) (Tzelepis et al. 2016), Avana (Project Achilles) (Meyers et al. 2017), and Jacquere (Drepanos et al. 2026) (**Figure 1b**). Each of these libraries have 4-5 sgRNAs target per gene with the total number of sgRNAs ranging from approximately 71,000 – 90,000 sgRNAs per library. These libraries each target 17,500 – 19,000 protein-coding genes. sgRNAs in the Yusa library consist of 19 nucleotide targeting sequence, while all other libraries have the 20-nucleotide design.

Previous surveys of multi-targeting and off-targeting sgRNAs have been conducted using the standard human reference genome 38 (GRCh38), and limited to the Avana and GeCKOv2 libraries (Aguirre et al. 2016; Schneider et al. 2017; Fortin et al. 2019). In this study, we use the T2T-CHM13 complete human reference genome (Nurk et al. 2022), which may detect additional off-target sites from previously uncharacterised genomic regions. Across the five libraries GuideRefine detected ∼2-6.7% of all sgRNAs as multi-targeting, ∼4-9.6% of sgRNAs as having single mismatch off-targets, and < 1% having PAM-distal double mismatches (**Figure 2a**). Aggregating multi-targeting and potential off-targeting sgRNAs, GuideRefine identified a large number of problematic sgRNAs in these libraries, ranging from 7.59% - 16.13% of sgRNAs.

**Figure 2.**
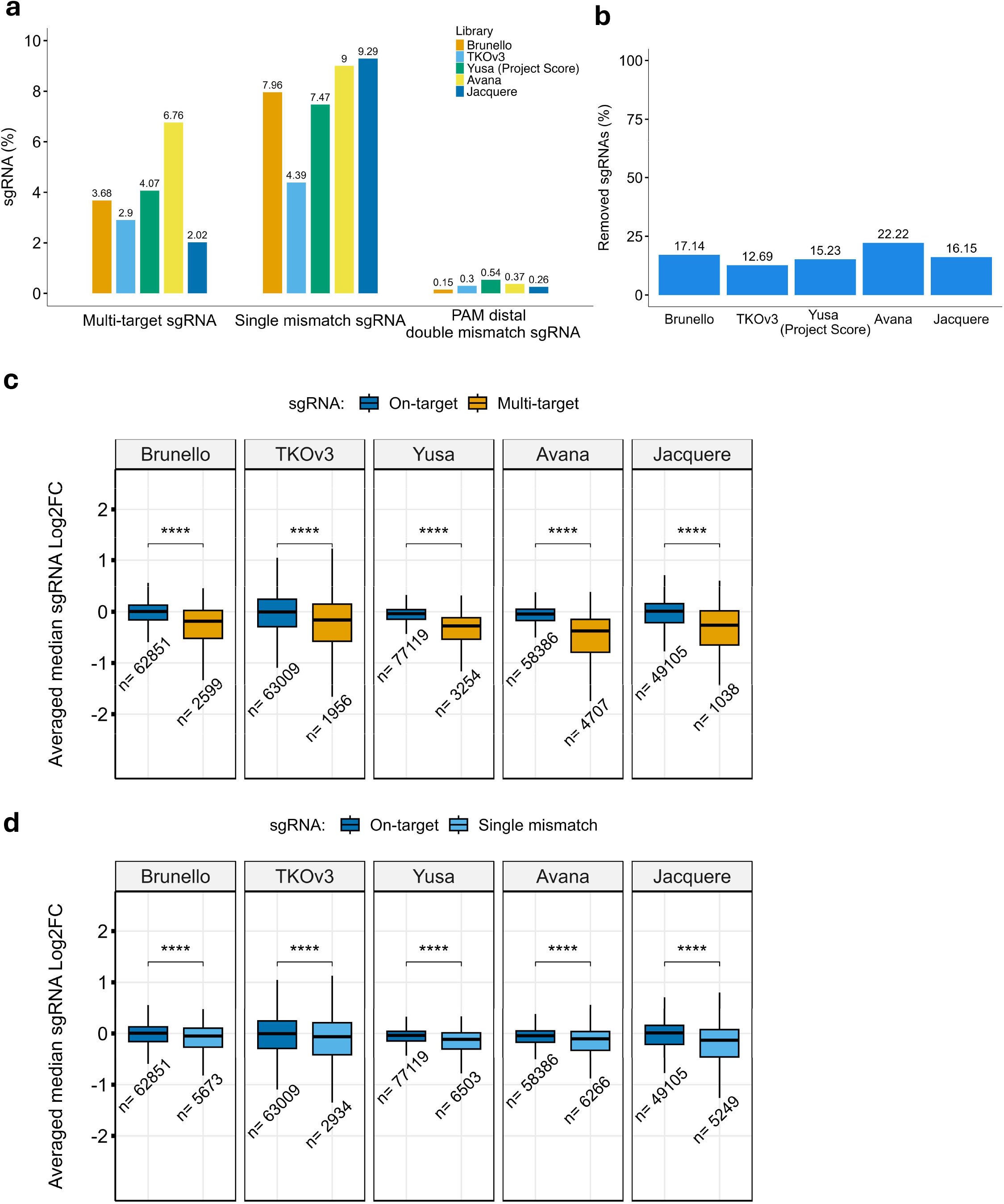
Multi-targeting and off-target sgRNAs are common across libraries and are associated with reduced cell fitness **(a)** A grouped bar chart showing the percentage of multi-targeting sgRNAs and off-targeting sgRNAs (single mismatch sgRNA, and PAM-distal double mismatch sgRNA) identified using T2T-CHM13 reference genome. **(b)** A bar chart showing the percentage of sgRNAs virtually removed across CRISPR-KO libraries **(c)** Boxplots showing the median sgRNA log-fold change between on-target and multi-targeting sgRNAs in all five libraries. **(d)** Boxplots showing the median sgRNA log-fold change between on-target and single-mismatch sgRNAs in all five libraries. For **(c)** and **(d)**, the horizontal thick line in the middle of the boxplot represents the median, the box edges represent the lower (Q1) and upper (Q3) quartiles, and the whiskers extend to values within 1.5 x interquartile range (IQR). All statistical tests were performed using Wilcoxon rank-sum; **** = *p* < 0.0001.

We then asked whether alignment of all sgRNAs against the T2T-CHM13 human reference genome can identify more problematic sgRNAs than the standard GRCh38 assembly. All problematic sgRNAs across five libraries were categorised based on the alignment against both genome builds. By using this approach, we uncovered less than 1% difference in the total number of problematic sgRNAs identified by using each of the genome builds (**Supplementary Figure 1a**), which suggests that the choice of genome assembly between GRCh38 and T2T-CHM13 reference genome has a negligible impact on the number of detected problematic sgRNAs. Therefore, unless otherwise stated, we discuss only the results with the T2T-CHM13 genome assembly for the remainder of the analysis. Following library refinement, GuideRefine virtually removed a total of 17.14% (Brunello), 12.69% (TKOv3), 15.23% (Yusa), 22.22% (Avana), and 16.15% (Jacquere) sgRNAs (**Figure 2b**). The differences in removal rates across libraries likely reflect their heterogeneous design strategies, including variations in the strategy to select high performing sgRNAs and the cell line contexts in which they were originally developed.

Multi-targeting sgRNAs present in the GeCKOv2 and Avana libraries have previously been shown to disproportionately reduce cell fitness, by causing excessive DNA double-strand breaks, through the joint disruption of two loci, or some combination of the two (Aguirre et al. 2016; Fortin et al. 2019). To verify if such findings extend to other libraries, we retrieved available sgRNA raw read count or log-fold change data from screens using each of the five libraries (**Figure 1b**). Where raw read counts were available, these were log-transformed and normalised to represent sgRNA log-fold change (see Methods). The log-fold change values for all sgRNAs were then integrated with their respective sgRNA alignment classification, thus categorising each sgRNA as either “on-target”, “multi-target”, and “single mismatch”.

We found that multi-targeting sgRNAs across all five CRISPR-KO libraries significantly impair cell fitness when compared to sgRNAs with only a single identified target locus (**Figure 2c**; Brunello *p* = 8.96×10^-203^, TKOv3 *p* = 4.74×10^-48^, Yusa *p* < 2.2×10^-16^, Avana *p* < 2.2×10^-16^, Jacquere *p* = 1.77×10^-85^; Wilcoxon rank-sum test). Furthermore, an increasing number of perfect on-target alignments was associated with progressively greater cell fitness defects, a trend that was consistent in all five libraries (**Supplementary Figure 1b–f**; Brunello *p* = 1.42×10^-206^, TKOv3 *p* = 1.25×10^-48^, Yusa *p* < 2.2×10^-16^, Avana *p* < 2.2×10^-16^, Jacquere *p* = 2.19×10^-86^; Jonckheere trend test). Altogether, these findings suggest that multi-targeting sgRNAs are detrimental to cell fitness, which is consistent in all five major CRISPR-KO libraries and with the observation previously reported by Fortin et al (2019) using the Avana library.

CRISPR-Cas9 DNA cutting has some degree of tolerance for imperfect base pairing, previous studies have established that single-nucleotide mismatches between sgRNA spacer and protospacer sequences can falsely direct Cas9 to unintended locations in the genome (Fu et al. 2013; Anderson et al. 2015; Zhang et al. 2015). Fortin et al (2019) demonstrated in the Avana library, that sgRNAs with a single-nucleotide mismatch occurring at the PAM-distal site are prone to off-target cleavage activity and contribute to a cell fitness defect. In our survey, single mismatch sgRNAs constitute the largest proportion of flagged problematic sgRNAs across the five CRISPR-KO libraries (**Figure 2a**). However, whether Cas9 driven by these sgRNAs can also cause cell fitness defects in the libraries included in this study, is currently unknown.

To address this, we applied the same strategy previously described, by using sgRNA sequences and log-fold change data to evaluate the effect of single mismatch sgRNAs on cell fitness. Single mismatch sgRNAs across all five CRISPR-KO libraries were found to impair cell fitness compared to sgRNAs without identified off-target matching (**Figure 2d**; Brunello *p* = 6.41×10^-59^, TKOv3 *p* = 5.19×10^-18^, Yusa *p* = 3.23×10^-183^, Avana *p* = 4.40×10^-81^, Jacquere *p* = 2.08×10^-165^; Wilcoxon rank-sum test). A higher number of single-mismatch off-target alignments was also associated with increasing cell fitness defects across all five libraries (**Supplementary Figure 1g–k**; Brunello *p* = 1.79×10^-61^, TKOv3 *p* = 2.63×10^-18^, Yusa *p* = 6.14×10^-169^, Avana *p* = 2.70×10^-85^, Jacquere *p* = 2.00×10^-169^; Jonckheere trend test). Taken together, these findings demonstrate that multi-targeting and single-mismatch off-targeting sgRNAs across major CRISPR-KO libraries can be detrimental to cell fitness and may confound CRISPR-KO screen analysis.

A survey of problematic sgRNAs across widely used libraries was recently reported by Drepanos et al. (2026) using CFD-based scoring (Drepanos et al. 2026). CFD is an off-target activity scoring metric based on mismatch tolerance at the sgRNA-DNA interface (Doench et al. 2016). CFD score ranges from 0 to 1, where higher scores indicate a greater probability for sgRNA off-target activity. As GuideRefine implements an alignment-based approach with pre-defined criteria rather than CFD-based scoring, we hypothesised that the two approaches would identify both overlapping and unique sets of problematic sgRNAs. To investigate this, we analysed the Jacquere library (4 sgRNAs/gene), the only library with fully annotated CFD scores for each sgRNA as reported by Drepanos et al. To ensure consistency between our analysis and that of Drepanos et al., GuideRefine was applied to the Jacquere library using the GRCh38 reference genome (see Methods). A comparison of problematic sgRNAs was conducted between 2,024 sgRNAs with CFD = 1.0 identified by Drepanos et al. and 8,449 problematic sgRNAs identified using GuideRefine (see Methods). Of the 2,024 sgRNAs flagged by Drepanos et al., 1,527 could be matched to a corresponding sgRNA in our GuideRefine classification. The remaining 497 sgRNAs were predominantly sgRNAs targeting non protein-coding genes defined by HGNC or lack canonical transcript with CDS in T2T-CHM13 GFF annotation, which were excluded from the working Jacquere library used in this study. This comparison revealed both overlapping and unique problematic sgRNAs between the two approaches (**Supplementary Figure 1l**), which reflects the fundamental differences in how sgRNA off-target activity is defined between the two approaches. Interestingly, all multi-targeting sgRNAs were concordantly identified by both approaches; however, this phenomenon did not extend to single mismatch and PAM-distal double mismatch sgRNAs.

We then focused our analysis on the single mismatch sgRNAs as a total of 6,807 single mismatch sgRNAs were not flagged by Drepanos et al., but detected by GuideRefine (**Supplementary Figure 1l**). We asked whether these set of sgRNAs could falsely direct Cas9 to off-target locations in the genome and disproportionally reduce cell fitness. To investigate this, we integrated the Jacquere sgRNA log-fold change data with our GRCh38 sgRNA alignment classification. These results were consistent with our T2T-CHM13 analysis; single mismatch sgRNAs were associated with significantly reduced cell fitness (**Supplementary Figure 1m;** *p* = 1.43×10^-174^; Jonckheere trend test). Interestingly, CFD scores confirmed that these sgRNAs had a median of maximum CFD score per sgRNA below 1.0 (median max CFD for sgRNAs with 0 single mismatch off-target alignment = 0 vs. median max CFD for sgRNAs with ≥ 1 single mismatch off-target alignment = 0.625) (**Supplementary Figure 1n and Supplementary Table 1a**). Altogether, these findings demonstrate that single mismatch sgRNAs, despite having a CFD score below 1.0, retain sufficient off-target activity to impair cell fitness and can be missed by a strict CFD score = 1.0 threshold alone.

### Genes consistently underrepresented in five libraries are characterised by short CDS length and limited PAM-sites

By removing problematic sgRNAs from the original pool, we then find that 12.73% (Brunello), 10.19% (TKOv3), 9.95% (Yusa), 16.29% (Avana), and 12.3% (Jacquere) of protein-coding genes are now targeted by fewer than three sgRNAs with no predicted off target effects (**Figure 3a**). We hypothesised that a set of genes may be consistently underrepresented across multiple libraries, reflecting the challenge in designing specific sgRNAs for those genes. To test this hypothesis, we compiled all genes with fewer than three sgRNAs across the five CRISPR-KO libraries and updated each gene symbol to ensure nomenclature consistency. We uncovered a total of 4,621 genes to be underrepresented, either uniquely in a single library or commonly across multiple libraries (**Figure 3b**). A total of 467 genes (∼10.10%) are consistently represented by fewer than 3 sgRNAs in each of the five libraries (**Supplementary Table 1b**), suggesting that these genes are inherently difficult to target with high specificity using current sgRNA design approaches.

**Figure 3.**
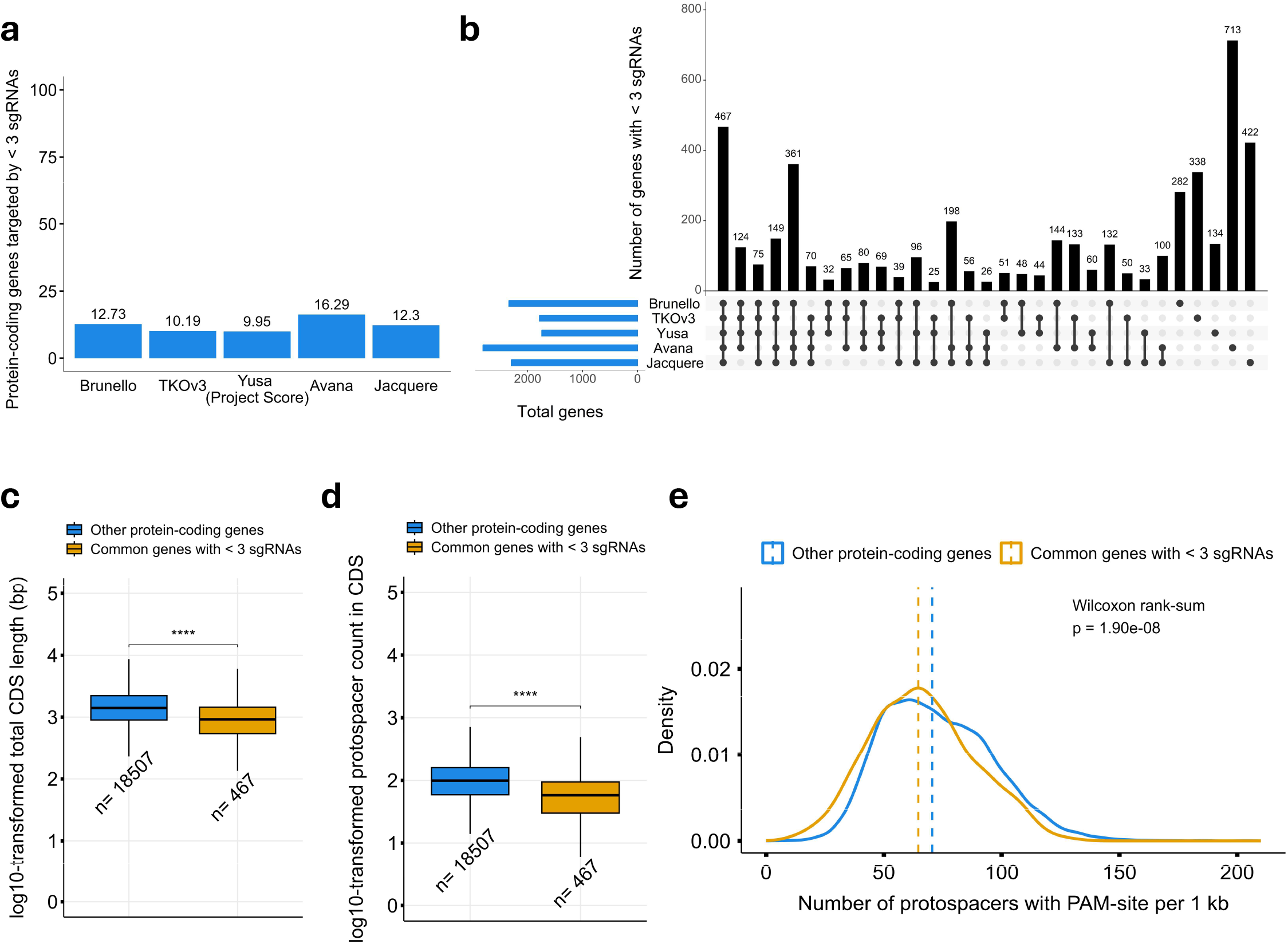
Protein-coding genes consistently underrepresented across all five libraries have shorter coding sequences and fewer PAM sites **(a)** A bar chart showing the percentage of protein-coding genes, as defined by HGNC, with fewer than three sgRNAs per gene. **(b)** An UpSet plot showing common and unique underrepresented genes across libraries **(c)** A boxplot representing log-transformed total CDS length between 467 consistently underrepresented genes and 18,507 other protein-coding genes **(d)** A boxplot showing the log-transformed protospacer count inside CDS region between 467 consistently underrepresented genes and other protein-coding genes. **(e)** The kernel density estimates of the number of protospacers with PAM-site per 1 kb, drawn using ggplot2 geom_density function. All statistical test were performed using Wilcoxon rank-sum; **** = *p* < 0.0001.

The plausible explanations for the complexity in targeting these genes are the short length of their CDS and limited number of available PAM sites. To understand whether the 467 genes have shorter CDS length on average compared to the rest of the protein-coding genes, we retrieved the total CDS length of all canonical and alternate transcripts for all protein-coding genes annotated in T2T-CHM13 reference genome (see Methods). We also mapped the remaining 18,507 protein-coding genes to T2T-CHM13 annotation to serve as the control comparison group. The 467 consistently underrepresented genes had significantly shorter CDS length on average compared to the rest of the protein-coding genes (**Figure 3c**; median CDS length in common genes < 3 sgRNAs = 912 base pairs vs. median CDS length in other protein-coding genes = 1,413 base pairs; *p* = 4.57×10^-36^, Wilcoxon rank-sum). This difference was further highlighted when the 467 genes were stratified by the CDS length, where 22.48% of these genes have less than 500 base pairs, compared to only 8.35% for the rest of the protein-coding genes (**Supplementary Figure 2a-b**).

We next sought to determine whether the shorter CDS lengths in all 467 genes were accompanied by a reduced number of available spacer sequences within their CDS region. We used *crisprDesign* to enumerate all possible spacer sequences with an NGG PAM site within the CDS of each gene (Hoberecht et al. 2022). The total number of possible spacer sequences was significantly lower in the 467 underrepresented genes compared to the rest of 18,507 protein-coding genes (**Figure 3d**; *p* = 8.45×10^-46^, Wilcoxon rank-sum). This reduction was further evident at the distribution level, where 42.83% of these genes have fewer than 50 spacer sequences in CDS region compared with only 19.08% of the rest of protein-coding genes (**Supplementary Figure 2c-d**).

To investigate whether the 467 consistently underrepresented genes also contained a reduced protospacer density, we calculated the number of protospacers with an NGG PAM site per kilobase of CDS for each gene. We discovered the 467 underrepresented genes had a significantly lower protospacer density compared to the rest of protein-coding genes (**Figure 3e**; median of common genes < 3 sgRNAs = 64.6 sites/kb vs. median of other protein-coding genes = 70.5 sites/kb; *p* = 1.90×10^-8^; Wilcoxon rank-sum). Altogether we demonstrated that the 467 consistently underrepresented genes across all five CRISPR-KO libraries are characterised by shorter CDS length, a smaller number of available spacer sequences, and importantly, a lower protospacer density within their coding regions. This set of protein-coding genes are more challenging to target with current sgRNA library design.

We then asked whether aggregating from multiple refined libraries the sgRNAs, which have no predicted multi-targeting or off-targeting activity, and applying sgRNA on-target efficacy scoring, could retain coverage of the 467 underrepresented genes. This could be useful for ensuring adequate representation of these genes in the design of future libraries. To investigate this hypothesis, we established a virtual mini-composite library containing all remaining sgRNAs targeting 467 genes which survived the library refining process (see Methods). Without additional filtering, 242 genes (∼51.8%) out of 467 consistently underrepresented genes could be targeted with at least three sgRNAs. To further select for sgRNAs predicted to direct Cas9 to cleave efficiently at their intended target, on-target activity scoring was applied to the aggregated sgRNAs using Rule Set 3, an on-target activity scoring metric that considers sgRNA sequence composition and small variations in the tracrRNA sequence (DeWeirdt et al. 2022). Rule Set 3 scores are continuous, where more positive scores indicate higher predicted sgRNA on-target activity and Cas9 cuts efficiently at the intended target location. By applying Rule Set 3 scores, the number of recoverable genes was reduced to 176 genes (∼37.7%) with at least three sgRNAs with high predicted Cas9 cutting activity (**Supplementary Table 1c**). These findings suggest that cross-library sgRNA aggregation can partially recover consistently underrepresented genes, and this approach can help improve the design of future CRISPR-KO libraries.

## DISCUSSION

CRISPR-KO screens are widely used to identify genetic vulnerabilities in many cancers, which supports drug discovery and target validation efforts. Several sgRNA libraries with varying design complexity are available for use in CRISPR-KO screens experiments. However, multi-targeting and off-targeting sgRNAs remain a persistent source of error in CRISPR-KO screen analysis. We have developed GuideRefine to help sgRNA library quality control in CRISPR-KO screen analysis. By virtually removing problematic sgRNAs from libraries, GuideRefine reduces the likelihood of false-positive gene hits and therefore, increases confidence in CRISPR-KO screen analysis that leads to a correct formulation of biological hypotheses. Beyond its primary function to refine sgRNA libraries, GuideRefine also generates an HTML report of library cleaning, an excel file containing gene-level sgRNA coverage after each refinement process, and a tabular file that contains all predicted problematic sgRNAs. These data outputs enabled us to perform a systematic survey of multi-targeting and off-targeting sgRNAs across five major CRISPR-KO libraries. In a broader context, these features facilitate the prospective identification of problematic sgRNAs in newly developed libraries and support the development of improved sgRNA design strategies.

GuideRefine has several limitations that should be noted. First, the stringent criteria of removing problematic sgRNAs can leave some genes with an insufficient number of sgRNAs for reliable gene-level statistical inference. Genes that are biologically relevant may fall below the minimum sgRNA threshold and thus be excluded entirely from the virtual refined library. A recent study proposed to mitigate this trade-off by balancing the composition of on-target and off-target sgRNAs in the library to avoid underrepresentation of genes in the library (Drepanos et al. 2026). In contrast, GuideRefine applies a conservative approach through a set of defined criteria for problematic sgRNAs. While this approach maximises sgRNA specificity, users should be aware of the potential for gene underrepresentation and are encouraged to examine the per-gene sgRNA coverage report generated by GuideRefine prior to continuing downstream analysis. Alternatively, users can disable the option to remove all single mismatch sgRNAs, given that our survey showed the majority of filtered sgRNAs across all five libraries belong to this category. This feature allows users to preserve greater sgRNA coverage as GuideRefine will only filter out multi-targeting sgRNAs, sgRNAs with single mismatches at PAM-distal positions, and sgRNAs with double mismatches at PAM-distal positions. Furthermore, some recently developed CRISPR-KO libraries were designed with only two sgRNAs per gene, such as MinLibCas9 (Gonçalves et al. 2021) and Gattinara library (DeWeirdt et al. 2020). These libraries are still compatible with GuideRefine, it can run the refinement process to completion and output the flagged problematic sgRNAs per gene, although all genes with fewer than three sgRNAs will be excluded in the sub-refined sgRNA library file. The major CRISPR-KO libraries contain a minimum of four sgRNAs target per gene and remain suitable with our pipeline. It should also be noted that GuideRefine does not assess sgRNAs based on Cas9 on-target cleavage activity, which is supported by other packages such as *crisprScore* (Hoberecht et al. 2022).

We found that the choice of genome assembly between GRCh38 and T2T-CHM13 has a negligible impact on the number of problematic sgRNAs identified across all five libraries. This suggests that despite T2T-CHM13 representing the first complete human genome assembly, the additional sequence content it harbours does not introduce a significant number of new multi-targeting or off-targeting sites for the sgRNAs present in these libraries.

We have shown that multi-targeting and single-mismatch sgRNAs across all five CRISPR-KO libraries are associated with reduced cell fitness. This confounding effect is likely driven by excessive DNA double-strand breaks induced at off-target genomic sites, which cause a cumulative DNA repair burden that compromises cell viability, a phenomenon that has been reported in the Avana library (Fortin et al. 2019), and the GeCKOv2 library (Aguirre et al. 2016). Additionally, Fortin et al. (2019) noted that sgRNAs co-targeting two coding regions, such as paralog pairs, can confound the interpretation of CRISPR-KO screen result. In our analysis we did not assess the effect of PAM-distal double mismatch sgRNAs on cell fitness due to the relatively small proportion of sgRNAs belong to this category in all five libraries (<1%), hence limiting the statistical power required for a meaningful comparison.

In this study we also compared problematic sgRNAs identified in the Jacquere library using GuideRefine’s alignment-based strategy with those identified by CFD off-target scoring conducted by Drepanos et al. (2026). This comparison revealed both overlapping and unique sets of problematic sgRNAs between the two approaches. We showed that a CFD score of 1.0 captures all multi-targeting sgRNAs but does not account for sgRNAs with lower probability of off-target activity, such as single mismatch sgRNAs. These results are likely due to the difference in both underlying methodologies, where GuideRefine identifies the problematic sgRNAs using a sequence-based alignment and a set of defined criteria (De Kegel and Ryan 2019; Fortin et al. 2019), while CFD scoring predicts the likelihood of off-target activity based on the position and identity of mismatches between sgRNA spacer and protospacer sequences (Doench et al. 2016). These two approaches capture partially overlapping but distinct sets of problematic sgRNAs. However, and importantly, we showed that the 6,807 single mismatch sgRNAs identified in Jacquere library but missed in Drepanos et al. at the CFD = 1.0 threshold, were on average associated with reduced cell fitness. Our findings highlight that GuideRefine alignment-based and pre-defined criteria strategy can capture biologically relevant off-target activity that CFD score thresholding alone may miss.

*In-silico* removal of problematic sgRNAs resulted in two categories of underrepresented genes: (1) genes that are underrepresented in some, but not in all five libraries, and (2) the 467 consistently underrepresented genes in all five libraries. For the first category, the aggregation of remaining sgRNAs from multiple libraries and followed by selection of sgRNAs based on predicted Cas9 cutting activity offers an immediate practical solution to resolve the sgRNA insufficiency issues. The cross-library sgRNA aggregation for these category of underrepresented genes also does not require de novo sgRNA design, a similar approach to that used in the development of the compact and optimised MinLibCas9 library, in which sgRNAs were compiled from Yusa, Avana, Brunello, and TKOv3 libraries before undergoing multiple steps of on-target and off-target scoring to select high-performing sgRNAs (Gonçalves et al. 2021).

For genes that belong to the second category, the challenge is more complex as there appear to be intrinsic genomic constraints in targeting them. In particular, their shorter CDS length and reduced NGG PAM site density fundamentally restrict the available sgRNA spacer design. We have shown that cross-library aggregation of remaining sgRNAs from four refined libraries, retaining only sgRNAs with a positive Rule Set 3 on-target activity score (Rule Set 3 > 0), can successfully recover 176 genes (∼37.7%). This relatively lenient on-target activity threshold was chosen given the severe sgRNA scarcity for these genes, as more stringent thresholds can substantially reduce the number of recoverable genes. Therefore, a trade-off exists between the number of sgRNAs with high on-target activity and the number of recoverable genes. While cross-library aggregation offers a practical temporary solution, a new sgRNA spacer design targeting the limited CDS regions of these genes will be required in the future library development.

For the remaining 291 genes (∼62.3%), cross-library aggregation returned fewer than three sgRNAs, further highlighting the severity of their genomic constraints. A comprehensive strategy to gain coverage for most of these genes would be to search for all possible sgRNA spacer sequences targeting their CDS regions using *crisprDesign* (Hoberecht et al. 2022). Then perform an *in-silico* removal of all problematic sgRNAs using GuideRefine, and followed by applying on-target activity scoring to select the top three best performing sgRNAs. This systematic methodology would likely recover a greater proportion of these 291 genes. To aid with the future library design, we provide comprehensive information of all 467 consistently underrepresented genes across five libraries in **Supplementary Table 1b**.

In this study, we presented GuideRefine, a computational pipeline for the detection and removal of problematic sgRNAs in CRISPR-KO libraries. Through a systematic survey of five widely used libraries, we showed the presence of multi-targeting and off-targeting sgRNAs and their detrimental impact on cell fitness. We also identified a set of protein-coding genes consistently underrepresented across all five libraries due to genomic constraints, representing a critical gap in the current library coverage. We anticipate that our pipeline and the surveys presented here will serve as a resource for the reinterpretation of existing CRISPR-KO screens and will aid in the development of next-generation sgRNA libraries with improved coverage, specificity, and biological interpretability.

## DATA AVAILABILITY STATEMENT

GuideRefine is available in GitHub (https://github.com/stefanusbernard/GuideRefine). Supplementary tables, materials, and analysis scripts used in this study are available in GitHub (https://github.com/stefanusbernard/CRISPR-KO-library-survey).

## ACKNOWLEDGMENTS

We are grateful to Dr. Metin Yazar and Dr. Olivier Dennler (C.J. Ryan group), and Dr. Aline Morrison, Dr. Pouya Abrari, Liadhan Farrell, and Julia Schymura (C. Santocanale group) for critical reading of the manuscript.

## SUPPLEMENTARY FIGURE LEGENDS

**Supplementary Figure 1.**
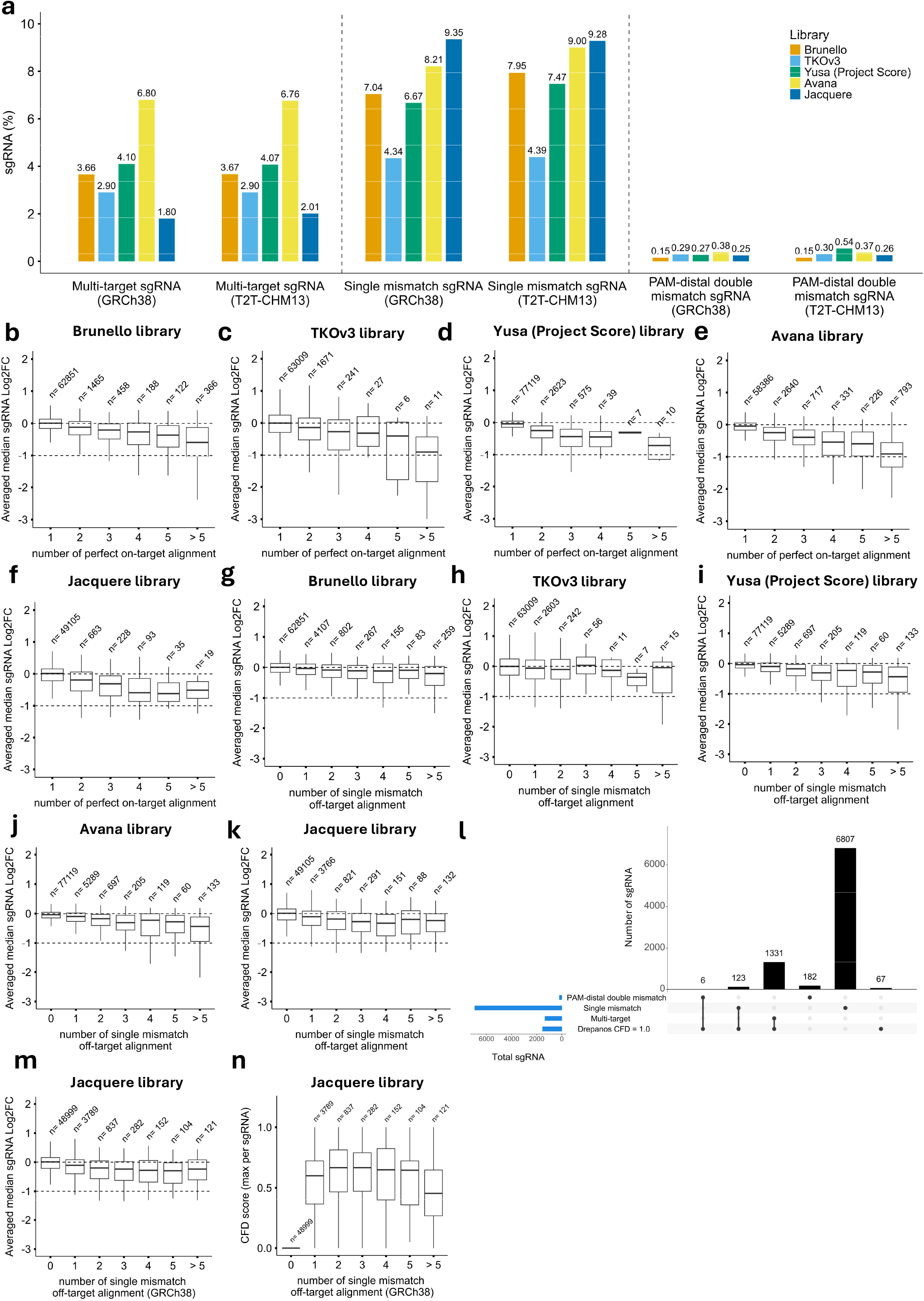
Multi-targeting and off-target sgRNAs are common across libraries and are associated with reduced cell fitness**. (a)** A grouped multi-bar chart showing the percentage difference of multi-targeting, single mismatch, and PAM-distal double mismatch sgRNAs between GRCh38 and T2T-CHM13 reference genome**. (b-f)** Stratified boxplots showing per-library sgRNA log-fold change across bins of increasing number of perfect on-target alignment **(g-k)** Stratified boxplots showing per-library sgRNA log-fold change distributions binned by single-mismatch off-target alignment count. **(j)** Problematic sgRNAs identified in Jacquere library by using GRCh38 reference genome between GuideRefine and Drepanos et al. CFD = 1.0. **(m)** Stratified boxplot showing sgRNA log-fold change across bins of increasing number of single mismatch off-target alignments. **(n)** Stratified boxplot showing maximum CFD score per sgRNA across bins of increasing number of single mismatch off-target alignments. For all panels using boxplot, the horizontal thick line in the middle of the boxplot represents the median, the box edges represent the lower (Q1) and upper (Q3) quartiles, and the whiskers extend to values within 1.5 x interquartile range (IQR). For **(m)** and **(n)**, sgRNAs were identified in the Jacquere library using GuideRefine with the GRCh38 reference genome

**Supplementary Figure 2.**
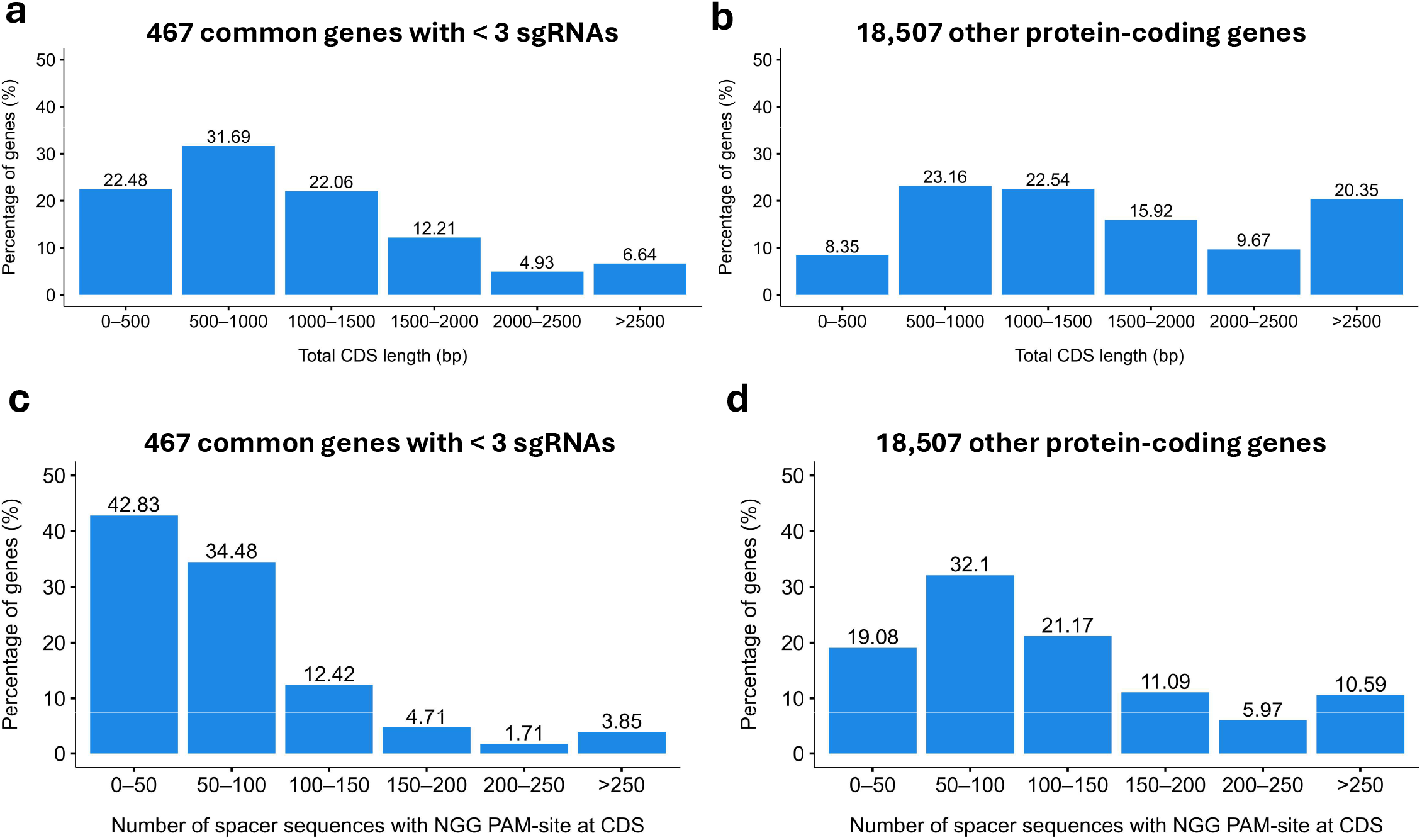
The 467 consistently underrepresented genes have shorter CDS length and fewer spacer sequences with NGG PAM sites at CDS. **(a-b)** Stratified bar charts showing the distribution of total CDS length between consistently underrepresented genes and the rest of protein-coding genes **(c-d)** Binned bar charts showing the distribution of total spacer sequences with NGG PAM-site at CDS region between consistently underrepresented genes and other protein-coding genes.

## SUPPLEMENTARY TABLE LEGENDS

**Supplementary Table 1a.** Median of CFD score of each bins of single mismatch off-target alignment in Jacquere library (GRCh38).

**Supplementary Table 1b.** The comprehensive information of 467 genes consistently underrepresented across five CRISPR-KO libraries after library refinement from multi-targeting and off-targeting sgRNAs.

**Supplementary Table 1c.** The virtual mini-composite library derived from cross-library on-target sgRNAs aggregation approach. This library comprises of 657 on-target sgRNAs targeting 176 consistently underrepresented genes.

## SUPPLEMENTARY DATA LEGENDS

**Supplementary Data 1.** sgRNA alignment classification using the T2T-CHM13 reference genome for the Brunello library

**Supplementary Data 2.** sgRNA alignment classification using the T2T-CHM13 reference genome for the TKOv3 library

**Supplementary Data 3.** sgRNA alignment classification using the T2T-CHM13 reference genome for the Yusa (Project Score) library

**Supplementary Data 4.** sgRNA alignment classification using the T2T-CHM13 reference genome for the Avana library

**Supplementary Data 5.** sgRNA alignment classification using the T2T-CHM13 reference genome for the Jacquere library

